# Hearing thresholds in the bat Carollia perspicillata vary by sex but not age

**DOI:** 10.1101/2024.10.21.619500

**Authors:** Olivia Molano, Emily Dale, Christine Portfors, Allison Coffin

## Abstract

Bats rely heavily on auditory perception for critical behaviors including navigation (echolocation) and conspecific communication. However, there is little information on the impacts of age and sex on the bat auditory system. Here, we used auditory brainstem response recordings to measure auditory thresholds in *Carollia perspicillata* (Seba’s short-tailed bat). We did not detect an age-related threshold shift at the age range examined (1-8 years old), suggesting that bats may be relatively resistant to age-related hearing loss. We had considerably more young bats (aged 1-2 years) than older animals (aged 4-8 years) so our results should be interpreted with caution. However, they are consistent with other studies showing that bats exhibit fewer signs of aging relative to other small mammals. In addition, we show significantly lower auditory thresholds among females compared to age-matched males, similar to sex differences reported in other mammals. Finally, we found increased thresholds in a single non-pigmented bat relative to age- and sex-matched conspecifics. While preliminary, this finding suggests that pigmentation may be important for inner ear function in bats, similar to results from other mammals.

## Introduction

In humans, age-related hearing loss, also called presbycusis, affects approximately 50% of those over age 75, with significant negative impacts on quality of life (Emmett and Francis, 2015; Goman and Lin, 2018). Presbycusis manifests as a progressive deterioration of hearing that typically begins at higher frequencies (Huang and Tang, 2010). While presbycusis occurs in both males and females, most studies report greater presbycusis in human males (Gates et al., 1990; Homans et al., 2017). Similarly, sex differences are reported in rodent models of both age-related and noise-induced hearing loss, as well as in physiological measures of hearing under baseline (undamaged) conditions (Kobrina and Dent, 2020; Milon et al., 2018; Shuster et al., 2019).

Presbycusis likely results from a combination of pathological features including damage to sensory hair cells and afferent neurons, as well as atrophy of the cochlear vasculature (Dubno et al., 2013; Pauler et al., 1988; Schuknecht and Gacek, 1993; Wu et al., 2020). Given the prevalence of this condition, there is a large body of research examining potential therapeutic solutions to prevent or reverse the impacts of age-related hearing loss (Bednar et al., 2015; Bowl and Dawson, 2019; Izumikawa et al., 2005; Watson et al., 2017; Zhao and Tian, 2022). To date, therapies rely on technological solutions such as hearing aids or - in extreme cases - cochlear implants; no biological therapies are yet available. Identifying a mammal that does not experience presbycusis may lead to a better understanding of the mechanisms underlying this form of hearing loss and provide therapeutic targets for disease prevention. One such possibility are echolocating bat species.

Many bat species use echolocation for navigation and foraging and therefore must depend on sensitive hearing throughout their lifespan (Galambos and Griffin, 1942; Griffin, 2001; Surlykke et al., 2014). Moreover, bats have unexpectedly long lives based on their body size (Brunet-Rossinni and Austad, 2004; Wilkinson and South, 2002), suggesting that they may retain their hearing abilities substantially longer than other small mammals. In this study, we examined hearing sensitivity of *Carollia perspicillata* (Seba’s short-tailed bat), a species that utilizes frequency-modulated echolocation calls and has a broad repertoire of social communication sounds (Knörnschild et al., 2013, 2014; Sterbing, 2002; Thies et al., 1998). In addition, *C. perspicillata* lives well in captivity and has an average lifespan of around 13 years, much longer than the lifespan of other small mammals such as the laboratory mouse (*Mus musculus*), which lives 3-4 years (Stewart et al., 2021). Given the strong reliance on hearing over their lifecycle, we hypothesized that *C. perspicillata* are resistant to age-related hearing loss. We also examined whether there are differences in hearing sensitivity of male and female bats, as has been demonstrated for many other mammalian species (Kobrina and Dent, 2020; Shuster et al., 2019).

## Methods

### Animals

We measured hearing sensitivity using the auditory brainstem response (ABR) in 25 *C. perspicillata* (16 females, 9 males) bats, with a body mass between 15-25 g. All bats were bred in captivity in a long-standing captive colony in the Portfors Lab bat facility at Washington State University Vancouver. All experiments were approved by the Institutional Animal Care and Use Committee at Washington State University (Protocol 6588).

### Auditory Physiology

We recorded ABRs to quantify hearing thresholds. Bats were anesthetized with subcutaneous injections of a cocktail containing 15-35 mg/kg ketamine, 8-15 mg/kg xylazine, and 1-1.5 mg/kg acepromazine (all drugs from Patterson Veterinary Supply), with juveniles requiring less anesthetic cocktail to obtain sufficient depth of anesthesia. Bats were placed on a heating pad kept at 24.5-26.5 °C inside a sound attenuation chamber and implanted with subdermal needle electrodes (CWE, Inc.), which were placed on the right mastoid next to the pinna, skull vertex over the midbrain, and mid-back along the spine (recording, reference, and ground electrodes, respectively). The active electrode was connected to the positive input of a Dagan 2400 differential amplifier. Signals were amplified by a factor of 2,000, bandpass filtered with a range of 100-3000 Hz (Khron-Hite 3550) and digitized at 25 kHz (National Instruments).

Pure tone stimuli were played through a calibrated leaf-tweeter speaker (LCY K100; Ying Tai) placed 10 cm away from the right ear and 45° away from the midline. We selected pure tones to allow us to compare our results with 88 other ABR studies in *C.perspicillata* (Capshaw et al., 2024a; Lattenkamp et al., 2021; Wetekam et al., 2020).The speaker was calibrated over a range of 3–100 kHz before each recording session using a microphone (model 4135, Brűel & Kjaer) positioned 10 cm from the speaker, where the animal’s right ear would be. Acoustic output was delivered via a 16-bit digital93 to-analog converter (500,000 samples/s, National Instruments) routed to a programmable attenuator (PA-5, Tucker Davis Technologies), then further routed to the speaker. Acoustic stimulation and data acquisition were controlled via custom software (Coffin et al., 2022; Felix et al., 2019).

We used pure tone stimuli (10, 20, 40, 60, 80 kHz, 5 ms duration, 1 ms rise/fall time, 10 rep/s), with alternating polarity to reduce stimulus artifact/cochlear microphonics in the final average waveform. Initial tones were played at 50 dB SPL (re: 20 μPa), with intensity initially adjusted down by 10 dB steps, then by 3 dB steps as the waveform diminished (Wetekam et al., 2020). Thresholds were based on the averaged response of 500 stimulus presentations per frequency and intensity level. Thresholds were visually determined based on the waveform, interpreted by two researchers, as seen in other studies on bat and rodent ABR responses (Capshaw et al., 2024a, 2024b; Felix et al., 2019; Uribe et al., 2023). Threshold was defined as the lowest intensity level at each frequency that yielded distinguishable peaks at wave 1 and wave 2/3, as show in Smotherman and Bakshi (2019) for the free-tailed bat *Tadarida brasiliensis*. Following ABR measurements, bats were given 5 mg of the anesthetic reversal agent Antisedan (Patterson Veterinary Supply). Bats recovered on a heating pad until they resumed normal activity, at which point they were returned to the bat facility.

### Statistical analysis

Data were analyzed using GraphPad Prism v. 10. ABR thresholds were first analyzed by two-way ANOVA using age or sex (Figs. 1 and 2, respectively) and frequency as the main effects, followed by post-hoc Bonferroni-corrected t-tests for multiple comparisons. In addition, threshold data were averaged across frequencies, then analyzed for an overall age effect using a Kuskal-Wallis non-parametric test (Fig. 1) or an overall sex effect using a Welch’s t-test (Fig. 2). Data are graphed as mean ± standard deviation.

**Fig 1.**
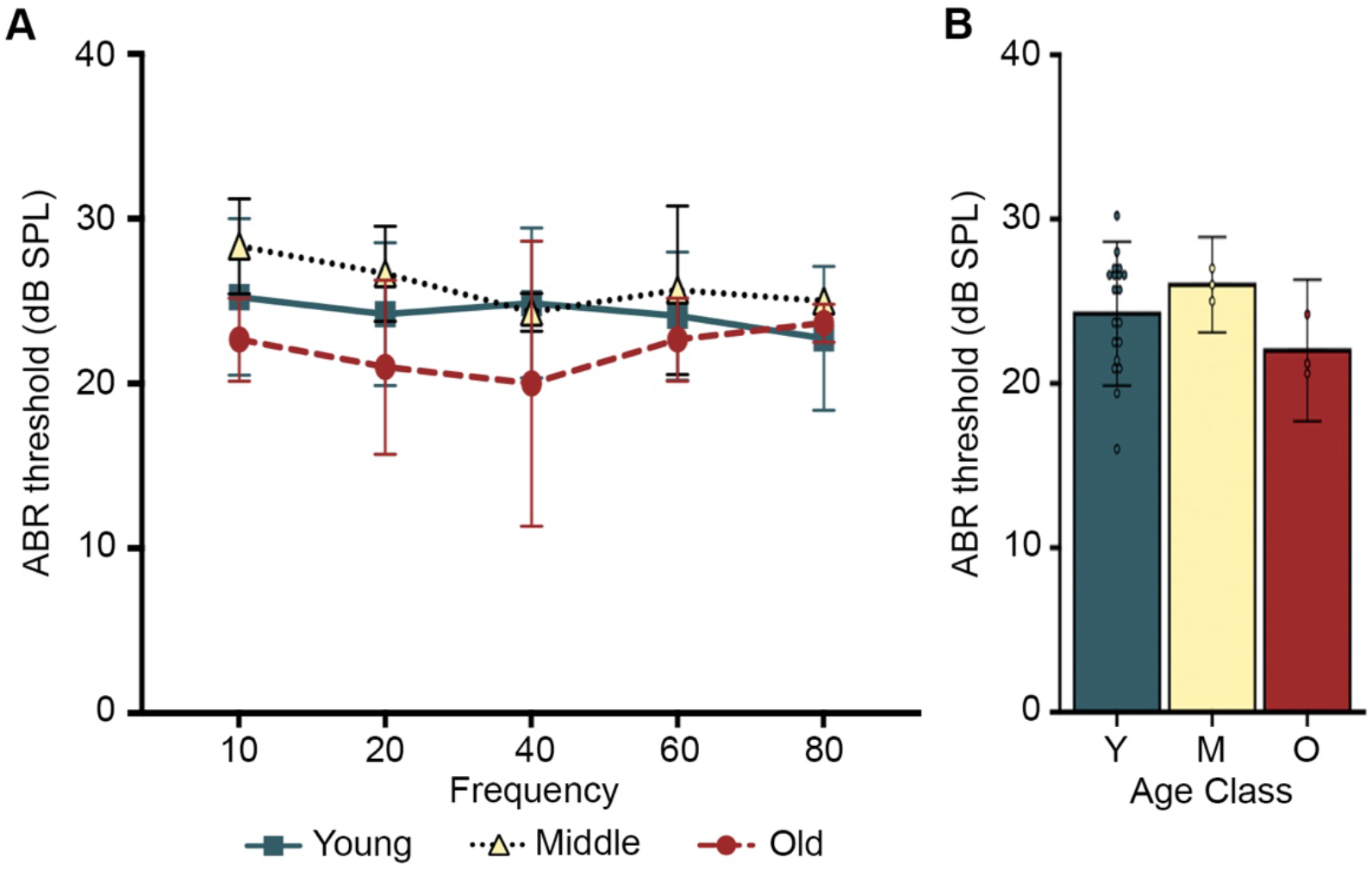
The effects of age on bat ABR thresholds. We grouped bats into three age classes, as described in Table 1. (A) There was a significant effect of age on frequency-specific thresholds [two-way ANOVA, main effect of age F_2,110_=3.245, p=0.043], with no effect of frequency [F_4,110_=0.486, p=0.746] or the interaction component [F_8,110_=0.440, p=0.895]. However, there were no significant differences at specific frequencies [Bonferroni-corrected t-tests]. (B) There was no significant age effect on overall threshold (summed across frequencies) [Krustal-Wallis non-parametric test, H=2.294, p=0.2318]. Both male and female animals were included in the experiment and pooled for analysis. N=19 Young, N=3 Middle Aged, N=3 Old. Data are presented as average ± 1 s.d. and dots in panel B represent individual bats.

**Fig 2.**
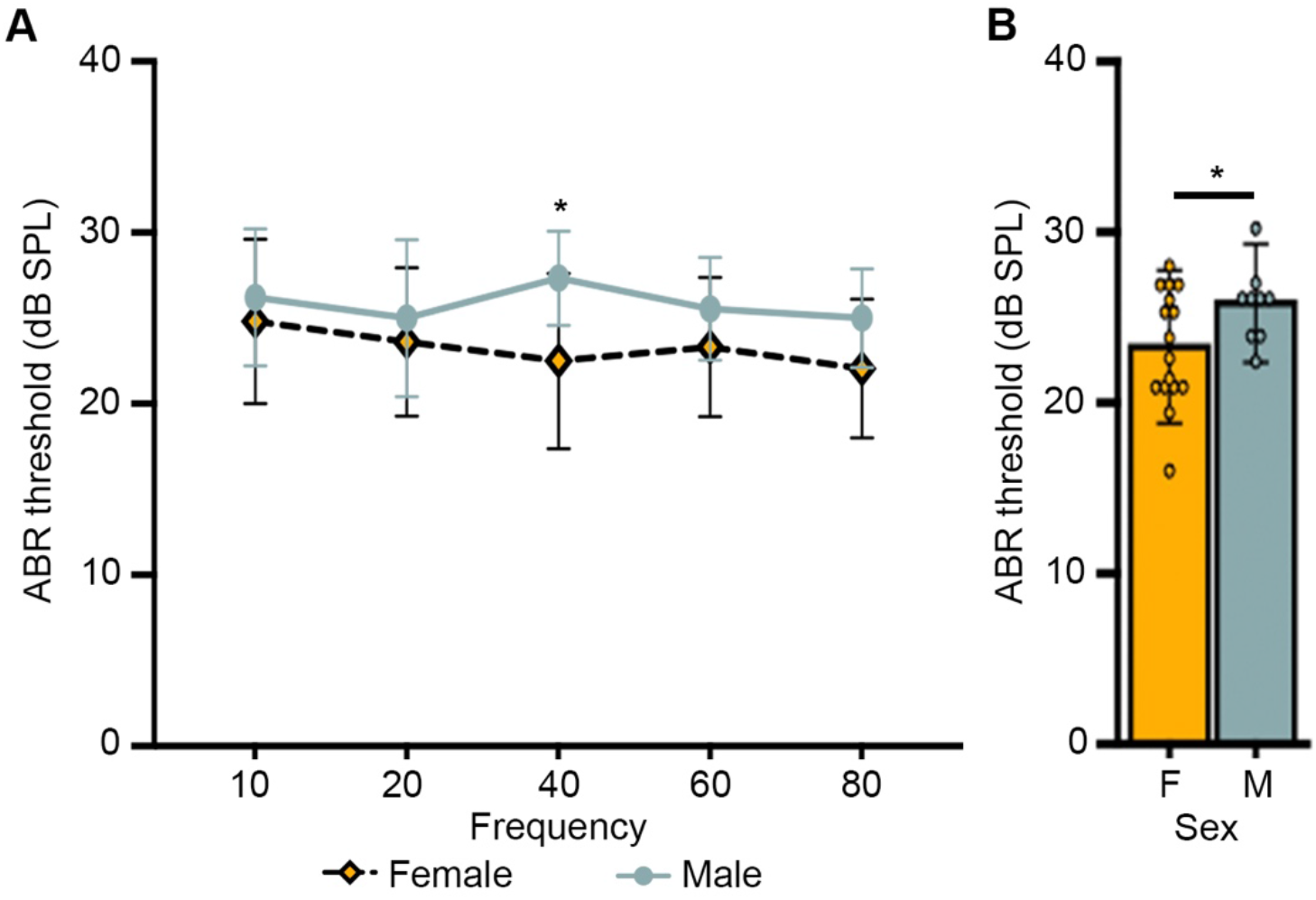
The effect of sex on bat auditory thresholds. (A) There was a significant effect of sex on frequency-specific thresholds [two-way ANOVA, main effect of sex F_1,115_=10.81, p=0.0013], but no effect of frequency [F_4,115_=0.718, p=0.581] or the interaction between main effects [F_4,115_=0.671, p=0.581]. Bonferroni-corrected t-tests show a significant threshold difference at 40 kHz [p=0.0321, indicated by *]. (B) There is an overall significant effect of sex when thresholds are averaged across frequencies [Welch’s t-test, p=0.0344]. N=16 Females, N=9 Males. Data are presented as average ± 1 s.d. and dots in panel B represent individual bats.

## Results

We first asked if there were age-related changes in ABR thresholds in *C. perspicillata* (Fig. 1). As shown in Table 1, we grouped individuals by their approximate ages, into young (1-2 years old), middle-aged (4 years old), and old (7-8 years old) categories.

**TABLE 1.** Sample sizes grouped by age.

| AGE GROUP (YEARS) | FEMALES | MALES | TOTAL |
| --- | --- | --- | --- |
| YOUNG (1-2) | 11 | 8 | 19 |
| MIDDLE (4) | 3 | 0 | 3 |
| OLD (7-8) | 2 | 1 | 3 |

We found a significant effect of age on auditory threshold (2-way ANOVA, main effect of age, p = 0.043) but upon post-hoc analysis did not see significant differences at any particular frequency (Fig. 1A). We therefore calculated the average threshold across frequencies for each age group. We observed no differences in average threshold (Fig. 1B); averaged thresholds were 23.2 ± 3.5 dB SPL, 26 ± 1 dB SPL, and 22.9 ± 1.9 dB SPL (mean ± s.d) for young, middle-aged, and old animals, respectively. These data suggest that hearing thresholds do not change with age in this species, at least not at the ages examined here. However, it’s important to note that we had relatively few animals for the middle-aged and old groups (N=3 per age class), so all conclusions are preliminary.

We next asked if ABR thresholds differed by sex (Fig. 2). We observed a significant sex difference (2-way ANOVA, main effect of sex, p=0.0013), with lower average thresholds in female bats at 40 kHz (Fig. 2A). Again, we then averaged across frequencies and confirmed our finding that females had lower ABR thresholds; females had averaged thresholds of 23.3 ± 3.4 dB SPL, while male thresholds were 25.8 ± 2.3 dB SPL (Fig.2B).

Finally, we took advantage of a spontaneously occurring pigment mutation that arose in our bat colony to ask if there was a threshold difference between a young albino male and his age- and sex-matched conspecifics. Interestingly, we observed higher thresholds in the albino male vs. pigmented 1-year-old male bats (Fig. 3). Unfortunately, we only had one non-pigmented animal and therefore could not conduct a statistical analysis. Further research is needed to investigate these trends with a larger data set from multiple non-pigmented bats and age- and sex-matched pigmented peers.

**Fig 3.**
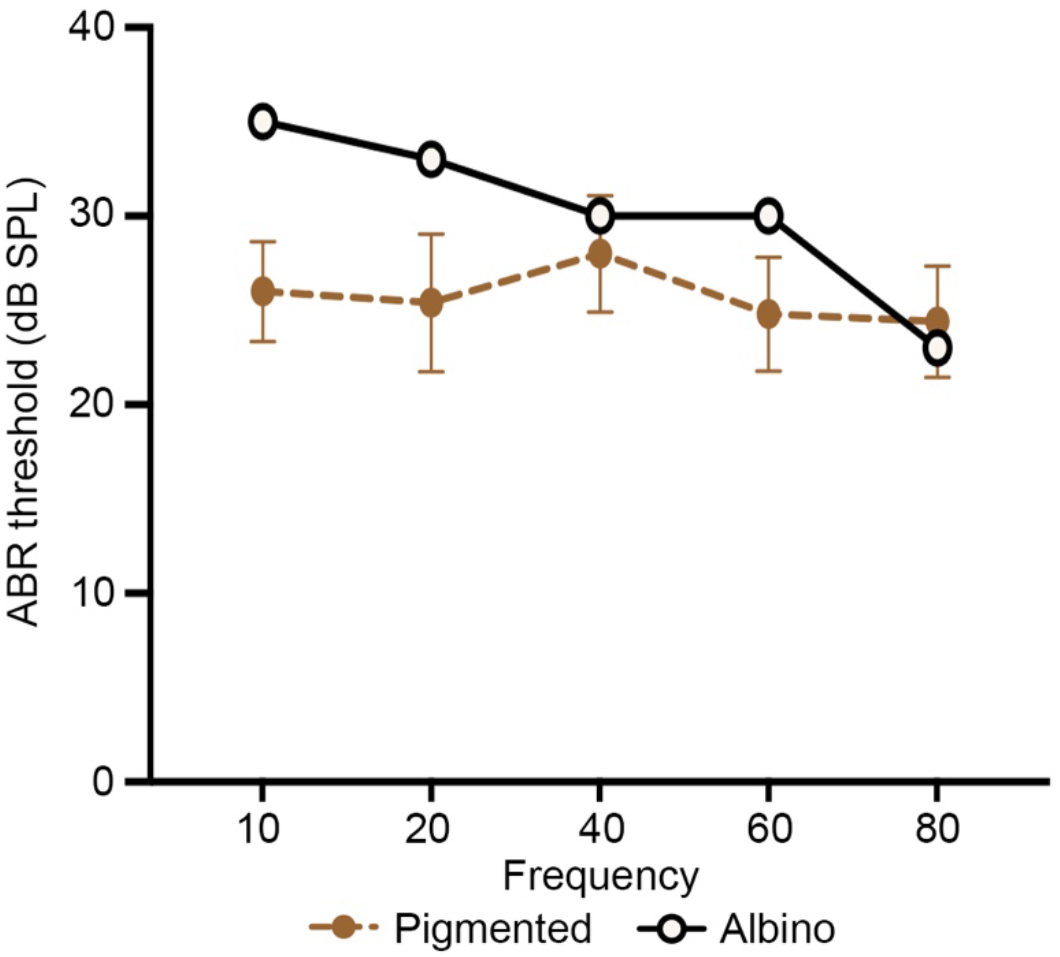
Average thresholds in pigmented vs. nonpigmented bats. Data show ABR thresholds obtained for the non-pigmented animal (open circles) vs. pigmented 1-year-old male bats (filled circles) (pigmented N=7 and nonpigmented N=1). Data are presented as average ± 1 s.d. for the pigmented bats.

## Discussion

Among mammals, bats are thought to have particularly acute hearing primarily because of their reliance on echolocation for prey capture and spatial orientation, but also due to their complex social communication sounds (Geipel et al., 2021; Ye and Luo, 2022). In this study, we used ABR measurements to quantify hearing thresholds in the echolocating fruit bat, *C. perspicillata*. An existing colony of diverse ages and both sexes presented the opportunity to examine threshold differences across age and sex.

Overall, we recorded ABR thresholds between 20-25 dB SPL over the frequency range of 10-80 kHz. These values are slightly higher than another ABR study in this bat species, which reported thresholds as low as 10 dB SPL in the most sensitive portion of the audiogram (20-60 kHz) (Wetekam et al., 2020). By contrast, Lattenkamp et al. (2021) report ABR thresholds closer to 50 dB SPL in *C. perspicillata* for the most sensitive frequencies tested (20-50 kHz), while Capshaw et al. (2024a) found ABR thresholds around 20-25 dB SPL over a portion of the frequency range (20-40 kHz), similar to our results. ABR thresholds can vary based on electrode placement, filtering techniques, and other conditions, including the threshold analysis metric selected (Wetekam et al., 2020; Schrode et al., 2022). The Wetekam et al. and Lattenkamp et al. studies determined threshold based on the root mean square of the waveform amplitude. Capshaw et al. (2024a) based their thresholds on observations by two independent observers, similar to our threshold determination method. Each study reports different thresholds, making it difficult to precisely compare between studies. However, within our population, we can confidently compare thresholds across sex and age.

We did not detect significant age-related hearing loss at the ages examined here. By contrast, a wealth of evidence shows robust age-related changes in the auditory system of humans and most other mammals (e.g., Gates et al., 1990; Dubno et al., 2013; Keithley, 2020). We are aware of only two other studies on presbycusis in bats, which show differing results from one another. Tarnovsky et al. (2023) demonstrated age-related hearing loss in wild-caught Egyptian fruit bats (*Rousettus aegyptacus*), with an average threshold loss of ∼1 dB SPL/year. However, using both ABRs and distortion product otoacoustic emissions, Capshaw et al. (2024b) found no age-related hearing loss in wild-caught big brown bats (*Eptesicus fuscus*). *R. aegyptacus* produces relatively simple echolocation calls using tongue clicks and relies on both hearing and vision for spatial navigation (Finger et al., 2023; Holland et al., 2004). By contrast, *E. fuscus, C. perspicillata* and other microchiropteran bats produce more complex echolocation vocalizations using the larynx and are generally considered to have more precise echolocation abilities (Fenton, 2003). Therefore, there may be additional selection pressure to preserve hearing in *C. perspicillata and E. fuscus* as compared to *R. aegyptacus*.

Resistance to auditory degradation may be a common feature of echolocating bats, with multiple studies demonstrating that bats are not susceptible to noise-induced hearing loss (Liu et al., 2021; Simmons et al., 2015, 2016; Simmons and Simmons, 2024). For example, a 1-hr exposure to a 116 dB SPL broadband noise did not alter behavioral thresholds in *E. fuscus*, nor did a more intense noise exposure (1 hr, 123 dB SPL) impact the ability of this species to use echolocation cues to navigate a cluttered environment (Simmons et al., 2016, 2018). An extensive comparative study by Liu et al. (2021) arrived at similar conclusions. Exposure to 120 dB SPL noise for 2 hr did not cause a threshold shift or hair cell loss in five different species of echolocating bats, while the non-echolocating fruit bat *Cynopterus sphynic* exhibited significant hearing loss from this stimulus (Liu et al., 2021). Collectively, these studies demonstrate that the inner ears of echolocating bats are remarkably resistant to both acute over-stimulation (noise) and age-related damage.

It should be noted, however, that *C. perspicillata* is known to live up to 13 years in captivity (Stewart et al., 2021), while the oldest bat included in the present study was 8 years old. Therefore, our “old” bats do not represent the maximum lifespan in this species. Additionally, our “old” cohort consisted of only 3 individuals (the total number available in our colony), so our data should be interpreted cautiously; additional studies are needed with a larger cohort of older bats. However, bats are known for their resilience to pathogenic insult and this ability to tolerate infection may be related to their unique inflammatory regulation (Ahn et al., 2023; Gorbunova et al., 2020). As inflammation is likely a major contributor to the development of age-related hearing loss (Kociszewska and Vlajkovic, 2022; Watson et al., 2017), it is likely that bats such as *C. perspicillata* are relatively resistant to presbycusis.

Next, we examined ABR thresholds from age-matched individuals of different sexes. Here we noticed significantly lower auditory thresholds among female individuals. This finding is unsurprising, as many mammals exhibit sex differences in auditory sensitivity, as can be seen from measurements of otoacoustic emissions and ABRs (Canlon and Frisina, 2009; Shuster et al., 2019). However, we did observe a large degree of variability between individuals, particularly for females, with threshold differences of up to 15-20 dB SPL at some frequencies. Our observation that female bats have higher auditory sensitivity than males is in line with other mammalian models (Canlon and Frisina, 2009; Shuster et al., 2019) and further emphasizes the point that using multiple animal models is beneficial to understand aspects of hearing such as sex-related differences in sound sensitivity.

These sex differences in ABR thresholds are unlikely to impact echolocation. In *C. perspicillata*, echolocation calls have an average level of 99 dB SPL at the source and echo return intensities above 50 dB SPL, with echo intensity increasing to as high as 80 dB SPL with decreasing distance between the bat and echo target (Brinkløv et al., 2011; Jakobsen et al., 2013; Beetz, 2017). In our study, ABR thresholds at high frequencies (those associated with echolocation) were 30 dB SPL or less, regardess of sex, meaning that slightly higher thresholds in males would not affect echo detection. In addition to echolocation, *C. perspicillata* also relies on acoustic cues for social communication. In this species, females respond to calls from vocalizing pups and male courtship vocalizations (Knörnschild et al., 2013, 2014). Therefore, lower thresholds may increase the ability of female bats to identify young or discriminate among mates. Future experiments with natural stimuli are necessary to test this hypothesis.

We had the unique opportunity to compare an albino individual’s auditory thresholds with pigmented, age- and sex-matched peers. With this investigation, we found higher thresholds in the albino individual, with the highest thresholds at the lowest frequencies tested (10-20 kHz). Therefore, the albino bat would likely have more difficulty detecting low-intensity social vocalizations but not echolocation signals, perhaps reflecting differential selection pressure on hearing associated with different tasks. Albinism is often co-morbid with sensory impairment, such as deafness associated with hypopigmentation in cats and guinea pigs and with oculocutaneous albinism in humans (Conlee et al., 1988; Dumitrescu et al., 2021; Lezirovitz et al., 2006; Strain, 2012). While this is a single case study and our result needs to be verified with a larger sample size, albino bats may present an additional model to study the intersection of pigmentation and hearing in an animal often considered a hearing specialist.

## Acknowledgments

This work was supported by funding from Washington State University Vancouver. We thank T. Hayward, A. Pederson, and N. Smith for assistance with animal care.

## Author Declarations

The authors declare that they have no conflicts to disclose.

## Data Availability

ABR data are available upon request.

